# Analysis of a mouse germ cell tumor model establishes pluripotency-associated miRNAs as conserved serum biomarkers for germ cell cancer detection

**DOI:** 10.1101/2023.09.09.556995

**Authors:** Amanda R. Loehr, Dennis M. Timmerman, Michelle Liu, Ad J.M. Gillis, Melia Matthews, Jordana C. Bloom, Peter K. Nicholls, David C. Page, Andrew D. Miller, Leendert H.J. Looijenga, Robert S. Weiss

## Abstract

Malignant testicular germ cells tumors (TGCTs) are the most common solid cancers in young men. Current TGCT diagnostics include conventional serum protein markers, but these lack the sensitivity and specificity to serve as accurate markers across all TGCT subtypes. MicroRNAs (miRNAs) are small non-coding regulatory RNAs and informative biomarkers for several diseases. In humans, miRNAs of the miR-371-373 cluster are detectable in the serum of patients with malignant TGCTs and outperform existing serum protein markers for both initial diagnosis and subsequent disease monitoring. We previously developed a genetically engineered mouse model featuring malignant mixed TGCTs consisting of pluripotent embryonal carcinoma (EC) and differentiated teratoma that, like the corresponding human malignancies, originate in utero and are highly chemosensitive. Here, we report that miRNAs in the mouse miR-290-295 cluster, homologs of the human miR-371-373 cluster, were detectable in serum from mice with malignant TGCTs but not from tumor-free control mice or mice with benign teratomas. miR-291-293 were expressed and secreted specifically by pluripotent EC cells, and expression was lost following differentiation induced by the drug thioridazine. Notably, miR-291-293 levels were significantly higher in the serum of pregnant dams carrying tumor-bearing fetuses compared to that of control dams. These findings reveal that expression of the miR-290-295 and miR-371-373 clusters in mice and humans, respectively, is a conserved feature of malignant TGCTs, further validating the mouse model as representative of the human disease. These data also highlight the potential of serum miR-371-373 assays to improve patient outcomes through early TGCT detection, possibly even prenatally.

## INTRODUCTION

Malignant testicular germ cell tumors (TGCTs) are the most common cancers diagnosed in adolescent and young adult men 15-39 years old in the US, and incidence has increased almost 40% in the last 50 years.^1^ TGCT patients have a 5-year overall survival rate of 95% due to the sensitivity of TGCTs to cisplatin-based chemotherapy. The WHO classifies TGCTs based on their distinct germ cell developmental origins and histological compositions.^2^ Type I TGCTs include teratomas and yolk sac tumors that occur in neonates and children, and Type III TGCTs are spermatocytic tumors that develop in older men. The most common TGCTs are the Type II TGCTs that arise after puberty, develop from germ cell neoplasia *in situ* (GCNIS), and are believed to originate from primordial germ cells (PGCs) that failed to differentiate during embryonic development.^3^ Of the Type II TGCTs, seminomas resemble embryonic germ cells, like the GCNIS lesions they develop from, while non-seminomas contain one or more other histological components including embryonal carcinoma (EC), yolk sac tumor, choriocarcinoma, and teratoma.^2^ EC consists of malignant pluripotent cells that are capable of self-renewal and differentiation into embryonic and extra-embryonic lineages, thus giving rise to the diverse histogenesis observed in many non-seminomas.^4–7^

To inform TGCT diagnosis and prognosis, clinicians use a set of imaging techniques and serum protein markers to detect TGCTs and monitor their response to treatment.^8^ Since the 1970s, α-fetoprotein, β-human chorionic gonadotropin, and lactate dehydrogenase have been used as serum biomarkers of TGCTs.^9,10^ However, these traditional biomarkers lack sensitivity and specificity. Only 50% of seminomas and 75% of non-seminomas show elevated levels of any one of the three serum proteins, and they can be elevated due to other disease processes.^11^ Thus, a better universal biomarker for TGCTs is needed to enable early detection, facilitate accurate diagnosis, and identify disease recurrence.

MicroRNAs (miRNAs) are small, noncoding RNAs that interact with target messenger RNAs (mRNAs) to regulate their stability and translation, and are vital for development and many other biological processes.^12^ Over a decade ago, miRNAs from the miR-371-373 cluster were found to be specifically expressed in malignant TGCTs but not in benign teratomas.^13,14^ Since then, these miRNAs have been found to significantly outperform traditional serum biomarkers in sensitivity and specificity for TGCT detection, both in the context of initial diagnosis as well as follow up. A recent study including 616 TGCT patients and 258 controls found that serum miR-371a-3p levels were sufficient to distinguish TGCT patients from male controls with 90.1% sensitivity and 94.0% specificity.^15^ Serum miR-371a-3p levels have also been found to correlate with primary tumor size and clinical stage, and they can be used to evaluate response to treatment, with levels decreasing after chemotherapy and orchiectomy.^15,16^ miR-371a-3p is also a good marker for detecting disease relapse. In a cohort of 33 patients with localized disease who underwent orchiectomy, all 10 patients who relapsed showed elevated miR-371a-3p levels at the time of recurrence, and these recurrences could be detected on average two months earlier using serum miR-371a-3p than with traditional clinical methods.^17^ miR-371a-3p expression is specific to malignant GCTs, as miR-371a-3p is undetectable in the serum of patients with testicular cancers of non-germ cell origin as it is in cancer-free controls.^18^ One limitation of the miR-371a-3p biomarker is that it is unable to detect pure teratomas, suggesting that it is expressed by undifferentiated cells within TGCTs but not following differentiation.^15^

The mouse miR-290-295 cluster is orthologous to the human miR-371-373 cluster, and the miRNAs in these clusters have conserved seed sequences.^19^ Additionally, miR-290-295 miRNAs are highly expressed in pluripotent mouse embryonic stem (ES) cells, just as miR-371-373 miRNAs are expressed in pluripotent human ES cells.^20,21^ In mouse ES cells, miR-290-295 have been implicated in pluripotency maintenance, proliferation, and suppression of apoptosis.^22–24^ They are the first miRNAs to be expressed *de novo* in the embryo—as early as the 2-cell stage—and are critical for embryonic development.^25,26^ The miR-290-295 miRNAs are also expressed by PGCs, the cells from which TGCTs originate, and are important for PGC migration during embryogenesis.^26,27^ Therefore, we hypothesized that miR-290-295 would serve as serum biomarkers for the presence of murine malignant GCTs as well.

The germ cell-specific *Pten* and *Kras* mutant (gPAK) mouse is the first genetically engineered mouse model of malignant TGCTs.^4^ These tumors resemble human mixed non-seminomas, containing EC that has tumor propagating and metastatic activity and also differentiates into teratoma components. gPAK tumors are similar to human TGCTs in that they arise from PGCs during embryonic development and are highly responsive to cisplatin-based chemotherapy.^4^ Here, we report that EC cells express and secrete miR-290-295, and serum miR-290-295 levels can be used to detect EC-containing murine TGCTs prior to birth. Additionally, we find that a differentiation-inducing experimental therapeutic, thioridazine (TR)^28^, can eliminate EC from gPAK tumors and concomitantly reduces miR-290-295 levels. These findings further validate the gPAK mouse as an accurate model of human TGCTs and highlight opportunities for pursuing the use of miR-371-373/miR-290-295 in the early detection of subclinical disease, elucidating the biological roles of these miRNAs in TGCT pathogenesis, and for testing new therapeutics in an animal model in which tumor progression can be easily monitored through serum miRNA levels.

## RESULTS

### Cultured murine EC cells express and secrete miR-291-293 and lose expression upon differentiation

We previously characterized three EC cell lines that were derived from *Pten^-/-^* malignant murine TGCTs with (EC14) or without (EC3, EC11) a *Kras^G12D^* activating mutation, and reported that these EC cultures can be differentiated with the drugs thioridazine (TR, giving rise to TR3, TR11, and TR14 cell lines) and salinomycin (SAL, giving rise to SAL11 and SAL14 cell lines).^28^ TR or SAL treatment induces changes in EC cell morphology from an ES cell-like to a fibroblast-like state, accompanied by a loss of pluripotency marker expression and a loss of tumorigenic potential.^28^ To determine if miR-290-295 cluster miRNAs were expressed by mouse EC cells, we analyzed the expression of miRNAs from this family in the EC cell lines and differentiated counterparts. In particular, levels of miR-291a-3p (homologous to human miR-372-3p and 373-3p) and miR-292-3p and miR-293 (homologous to different isoforms of human miR-371a-3p) were measured as representative miRNAs in this cluster and collectively referred to as miR-291-293 hereafter (Supplemental Figure 1).^19^ miR-291-293 miRNAs were highly expressed by all three EC cell lines, and this expression was significantly reduced in the TR and SAL-differentiated derivatives (Figure 1A). As a precursor to determining if miR-291-293 were present in the serum of mice with TGCTs, we next tested if these miRNAs were secreted by cultured EC cells. miR-291-293 were detected in the conditioned media of all three EC cell lines, while very low to no expression was detected in conditioned media from the differentiated cells (Figure 1B). These findings suggest that miR-291-293 expression and secretion is specific to the pluripotent state of EC cells and significantly diminished upon differentiation.

**Figure 1.**
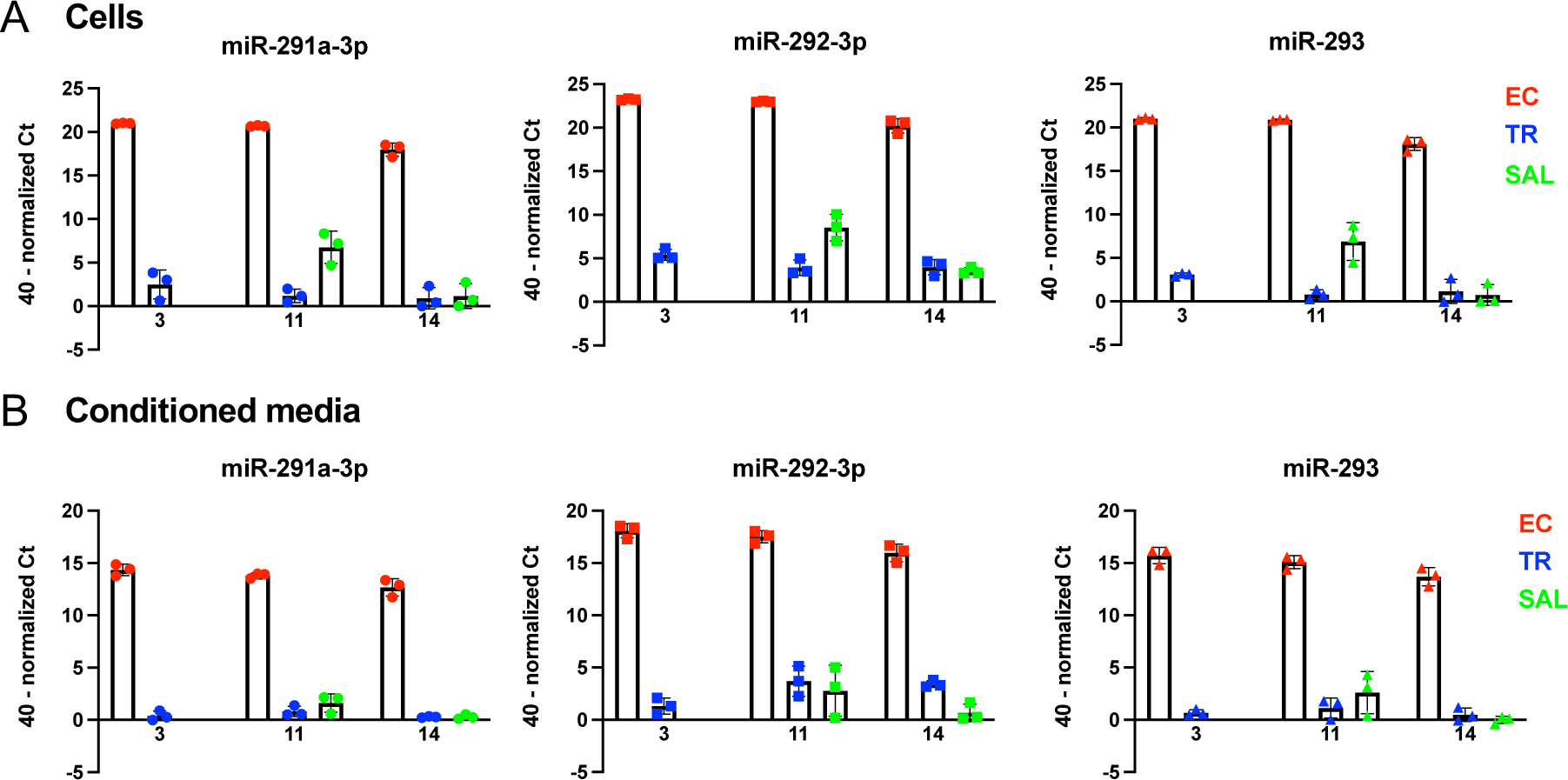
miR-291-293 expression in three independent EC cell lines (3, 11, and 14) is lost following thioridazine (TR)- or salinomycin (SAL)-mediated differentiation. miR-291-293 expression in (A) cells or (B) conditioned media. n = 3 independent replicates per group. Data are mean ± SD. Differences are significant for all comparisons between EC and TR or SAL groups (p < 0.001, multiple unpaired t-tests with Holm-Šídák correction for multiple comparisons).

### Predicted miR-290-295 targets are enriched in thioridazine-differentiated cells

To understand if miR-290-295 have a functional role in regulating gene expression in EC cells, we completed RNA sequencing (RNA-Seq) of all three EC cell lines and their TR-differentiated derivatives (part of this RNA-Seq data set was previously published^28^) and performed Gene Set Enrichment Analysis (GSEA)^29^ for predicted targets of mouse miRNAs. If miR-290-295 were negatively regulating their target genes, as miRNAs are canonically known to do, then the amount of RNA for their predicted target genes was expected to be enriched in TR-differentiated cells, since these cells lose miR-291-293 expression. The miRDB gene sets deposited in the Molecular Signatures Database consist of a gene set for each mouse miRNA catalogued in miRDB v6.0 containing high-confidence computationally predicted target genes as determined by the MirTarget algorithm.^30,31^ GSEA of the 1,769 miRDB gene sets revealed that the gene set consisting of predicted targets of miR-291a-3p and miR-294-3p (MIR_291A_3P_MIR_294_3P) was among the most highly enriched gene sets in TR-differentiated cells compared to EC cells for all three cells lines (17^th^ most highly enriched in TR3 cells, 14^th^ in TR11 cells, 26^th^ in TR14 cells) (Figure 2A). In fact, gene sets of predicted targets of many miR-290-295 miRNAs were significantly enriched in all three TR-differentiated cell lines compared to their parental EC cell lines (Figure 2B).

**Figure 2.**
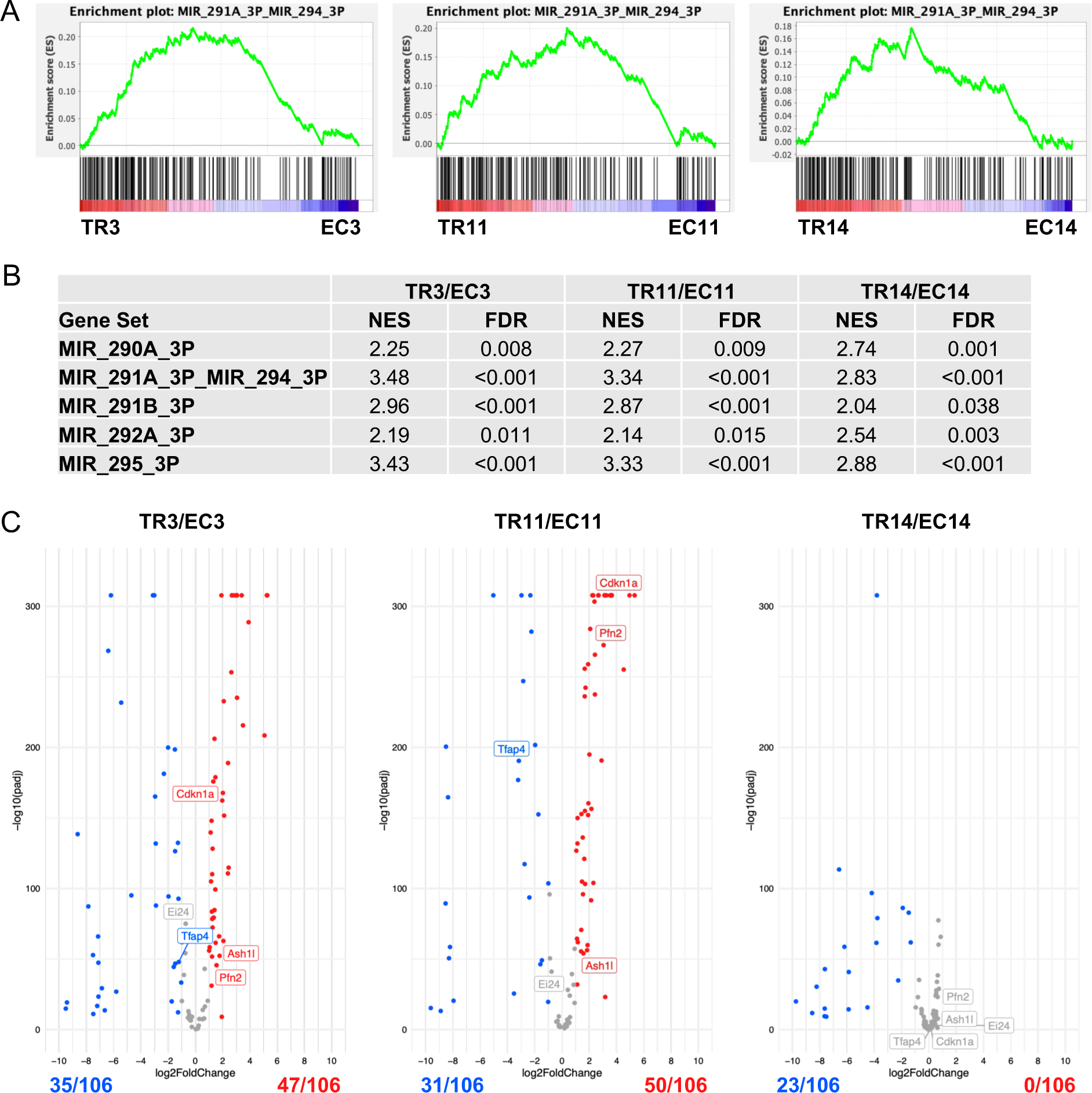
miR-290-295 targets are upregulated in TR-differentiated cells. (**A**) GSEA enrichment plots for the MIR_291A_3P_MIR_294_3P gene set across pre-ranked gene lists comparing gene expression in TR-differentiated cells to parental EC cell lines. (**B**) Significantly enriched (FDR < 0.05) mirDB gene sets associated with the miR-290-295 cluster in TR-differentiated cells compared to parental EC cell lines with normalized enrichment scores (NES) as determined by GSEA. (**C**) Volcano plots showing differential gene expression of the 106 candidate target genes of the miR-290-295 cluster from Schaefer et al.^40^ Gene names of experimentally determined direct targets of the miR-290-295 cluster are labelled. Genes that were significantly upregulated in TR-differentiated cells are in red, genes that were significantly downregulated in TR-differentiated cells are in blue, and genes that were not significantly differentially expressed are in grey. Significance is defined as a p adjusted value (padj) < 0.05.

A literature search was also undertaken to identify experimentally proven direct targets of the miR-290-295 family. In total, 26 target genes were identified from 16 publications, which were focused primarily on ES cells (Supplemental Table 1).^22–24,32–44^ Of these 26 genes, 13 were significantly upregulated and five were significantly downregulated in TR3 cells compared to EC3 cells (Supplemental Figure 2). Similarly, 14 were significantly upregulated and five were significantly downregulated in TR11 cells compared to EC11 cells. This suggests that miR-290-295 downregulate the mRNA levels of their target genes in EC cells, because expression of many target genes was increased following loss of miR-290-295 expression upon differentiation. The decreased RNA levels for some target genes following thioridazine-induced differentiation likely reflects the predominant influence of regulatory mechanisms other than loss of miR-290-295 expression. The limited changes in gene expression observed in EC14 cells after differentiation, with none of the known miR-290-295 target genes being significantly upregulated in TR14 cells, and only two being significantly downregulated, is discussed further below. Of the genes that were significantly upregulated in TR3 and TR11 cells compared to their parental EC cell line, *Cdkn1a, Rbl2,* and *Lats2* are all negative regulators of the G_1_/S transition that function to slow cell cycle progression.^23^

Schaefer *et al*., recently identified 360 genes predicted to be targets of the mouse miR-290-295 cluster.^40^ The target genes were identified through an integrative analysis of multiple datasets including computationally-predicted miR-290-295 targets in TargetScan, AGO2-miRNA binding and loading data from ES cells, and genes upregulated in global miRNA KO ES cells. 106 of these 360 candidate target genes were found to be upregulated following miR-290-295 KO in ES cells.^40^ Although miRNAs can also act by inhibiting the translation of their mRNA targets,^45^ this target gene set focuses on those mRNAs degraded by the miR-290-295 cluster. We queried the 106 gene targets of miR-290-295 in our RNA-seq data from EC and TR-differentiated cells and found that 47 of them were significantly upregulated and 35 were significantly downregulated in TR3 cells compared to EC3 cells (Figure 2C). Additionally, 50 genes were significantly upregulated and 31 were significantly downregulated in TR11 cells compared to EC11 cells. Together, these data indicate that many miR-290-295 targets are upregulated following EC cell differentiation, although several targets were significantly downregulated, likely because of large-scale reprogramming of gene expression following differentiation that is in part independent of miR-290-295. None of the 106 target genes were significantly upregulated and 23 were significantly downregulated in TR14 cells compared to EC14 cells. This may reflect the fact that, unlike TR3 and TR11, which only harbor a *Pten* inactivating mutation, TR14 cells, which have both *Pten* inactivating and *Kras* activating mutations, remain more highly proliferative and retain some transformed features following differentiation,^28^ with miR-290-295 target genes perhaps correspondingly less affected than in TR3 and TR11 cells. A greater extent of spontaneous differentiation in EC14 cultures as compared to EC3 or EC11 cultures also may obscure identification of differentially expressed genes following TR treatment of EC14 cells.

### Serum miR-291-293 detection is specific to mice with EC-containing tumors

We previously demonstrated that EC cell tumorigenicity is significantly reduced or fully abrogated after TR or SAL-mediated differentiation.^28^ Upon subcutaneous injection into immunocompromised host mice, all three EC cell lines formed tumors with EC and teratoma components. By contrast, the differentiated derivatives of EC3 and EC11 did not form tumors, and the differentiated derivatives of EC14 formed small sarcomas that grew much slower than the tumors from the parental EC14 cell line.^28^ For the present study, we tested whether miR-291-293 could be detected in the serum of mice bearing EC cell-derived tumors. Serum miR-291-293 levels were high in mice that developed tumors from the EC cell lines, but low to undetectable in mice that received differentiated cells (except for two mice that received TR11 cells and had detectable serum miR-291-293 levels despite having no identifiable tumor) (Figure 3A). One mouse that received EC3 cells and had very low levels of serum miR-291-293 formed only one small tumor in which no OCT4 staining was detected, indicating that the tumor may have lacked an EC component. Serum miR-291-293 levels were also low or undetectable in *Dazl^-/-^* mice with testicular teratomas, as well as in wild-type 129S6 male mice (Figure 3A). These data suggest that it is EC within murine TGCTs that expresses and secretes miR-291-293.

**Figure 3.**
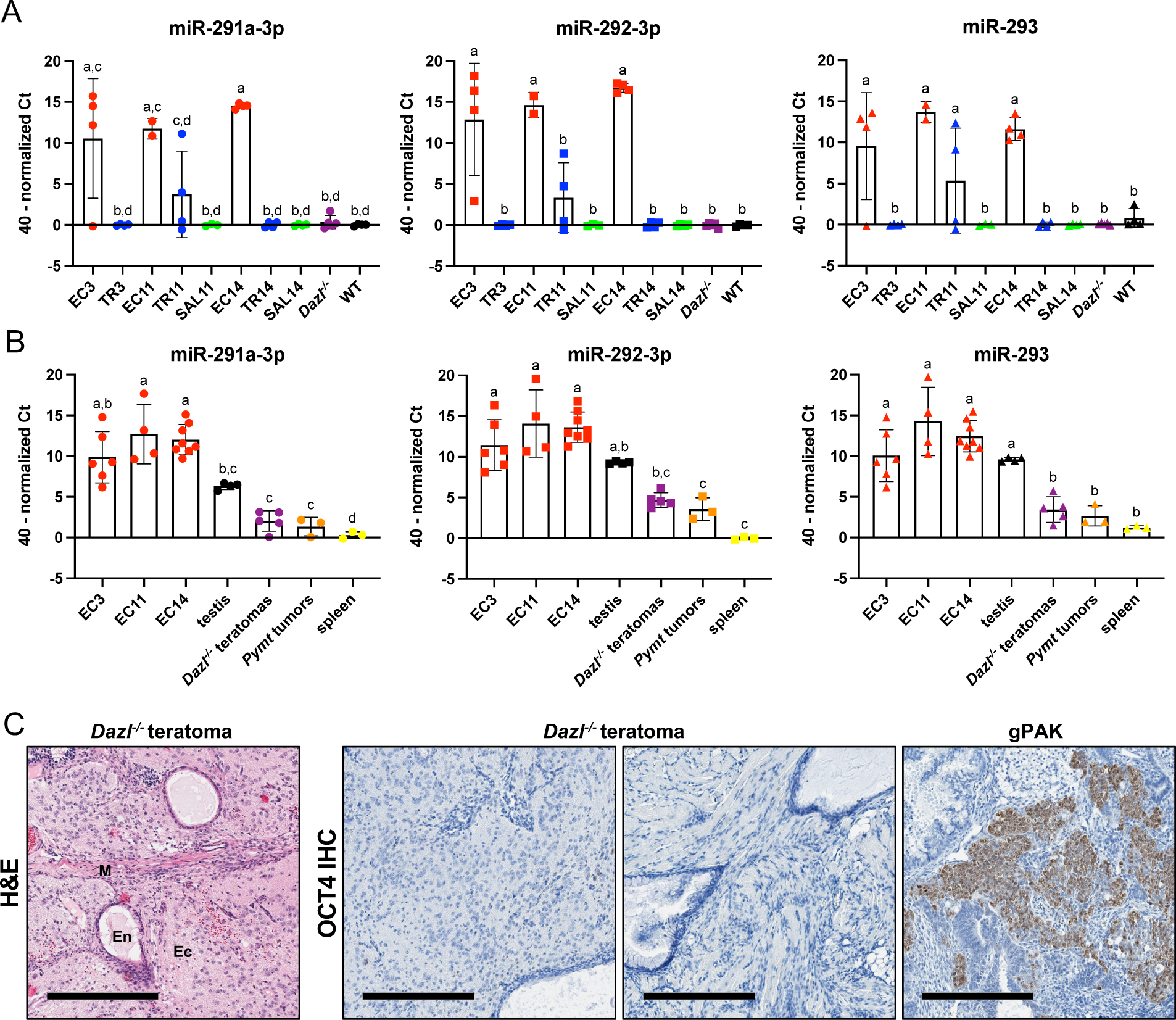
miR-291-293 expression is specific to EC-containing tumors. (**A**) miR-291-293 expression in serum for mice 2-3 weeks after receiving a subcutaneous injection of EC cells, TR- or SAL-differentiated cells, *Dazl^-/-^* mice with testicular teratomas, or wild-type 129S6 adult male mice. (**B**) miR-291-293 expression in EC-derived tumors, normal adult testis from 129S6 mice, teratomas from *Dazl^-/-^* mice, mammary adenocarcinomas from *Pymt* mice, or spleens from wild-type 129S6 mice. Data represented as mean ± SD. Significant differences exist between groups that do not share a letter (p < 0.05, ordinary one-way ANOVA and Tukey’s multiple comparisons test). (**C**) Representative image of H&E-stained *Dazl^-/-^*testicular teratomas with tissues derived from endoderm (En), mesoderm (M), and ectoderm (Ec) indicated and representative images of OCT4-IHC stained *Dazl^-/-^* testicular teratomas (n = 5) and a gPAK tumor (positive control). Scale bar = 300 μm.

miR-291-293 were also detectable in RNA extracted from EC-cell derived tumor tissue as well as normal adult testis tissue (Figure 3B). Expression in normal testis tissue was expected, as miR-290-295 and miR-371a-3p were previously shown to be expressed in mouse spermatogonia and normal human testis tissue, respectively.^27,46^ To confirm that miR-291-293 expression was specific to germ cells and EC cells, miR-291-293 levels were also measured in *Dazl^-/-^* testicular teratomas, MMTV-*PyMT* mammary adenocarcinomas, and normal adult mouse spleen. All three of these tissue types had significantly lower levels of miR-291-293 compared to EC-derived tumors (Figure 3B). H&E staining and OCT4 immunohistochemistry confirmed that, as previously reported^47^, the testicular tumors in adult *Dazl^-/-^* mice were pure teratomas containing tissues derived from all three embryonic germ layers, and unlike gPAK malignant teratocarcinomas, contained no detectable OCT4-positive EC (n = 5) (Figure 3C). These results suggest that miR-291-293 detection is specific to the presence of EC in the mixed TGCTs of gPAK mice. Interestingly, despite comparable levels of miR-291-293 in normal adult testis tissue and EC-containing tumors, the miRNAs were not detected in the serum of wild-type male mice (Figure 3A and B), observations consistent with previous findings in humans^46^. These results suggest that EC, but not adult germ cells, can secrete detectable levels of the pluripotency-associated miRNAs into circulation.

### Thioridazine treatment eliminates EC cells from gPAK tumors, reducing serum miR-291-293 levels

We next sought to determine if serum miR-291-293 levels were associated with the presence of EC cells in spontaneously occurring TGCTs in the gPAK mouse model and if expression would be lost following therapy-induced EC ablation *in vivo*. gPAK mice were treated with TR or vehicle control once every three days for three weeks starting at 21 days of age, when testicular tumors often can be first detected by palpation in this model.^4^ We previously found that TR treatment significantly extends the survival of mice with human EC cell xenografts, and in a transformed induced pluripotent stem (iPS) cell allograft model, TR treatment significantly reduces the percentage of OCT4-positive EC cells within tumors.^28^ Here, TR extended the survival of gPAK mice, although the effect was not statistically significant (Figure 4A). Of note, two of eight TR-treated mice reached the pre-defined endpoint of 70 days, whereas only one of the nine control mice lived past 42 days and none lived beyond 61 days. To determine EC cell abundance within tumors from control and TR-treated mice, we performed OCT4 immunohistochemistry and quantified the percentage of tumor area that was OCT4-positive. Except for one TR-treated mouse that had two tumors with OCT4-positive cells, tumors from TR-treated mice were depleted of OCT4-positive cells, whereas five of nine control tumors contained OCT4-positive cells (Figure 4B&C).

**Figure 4.**
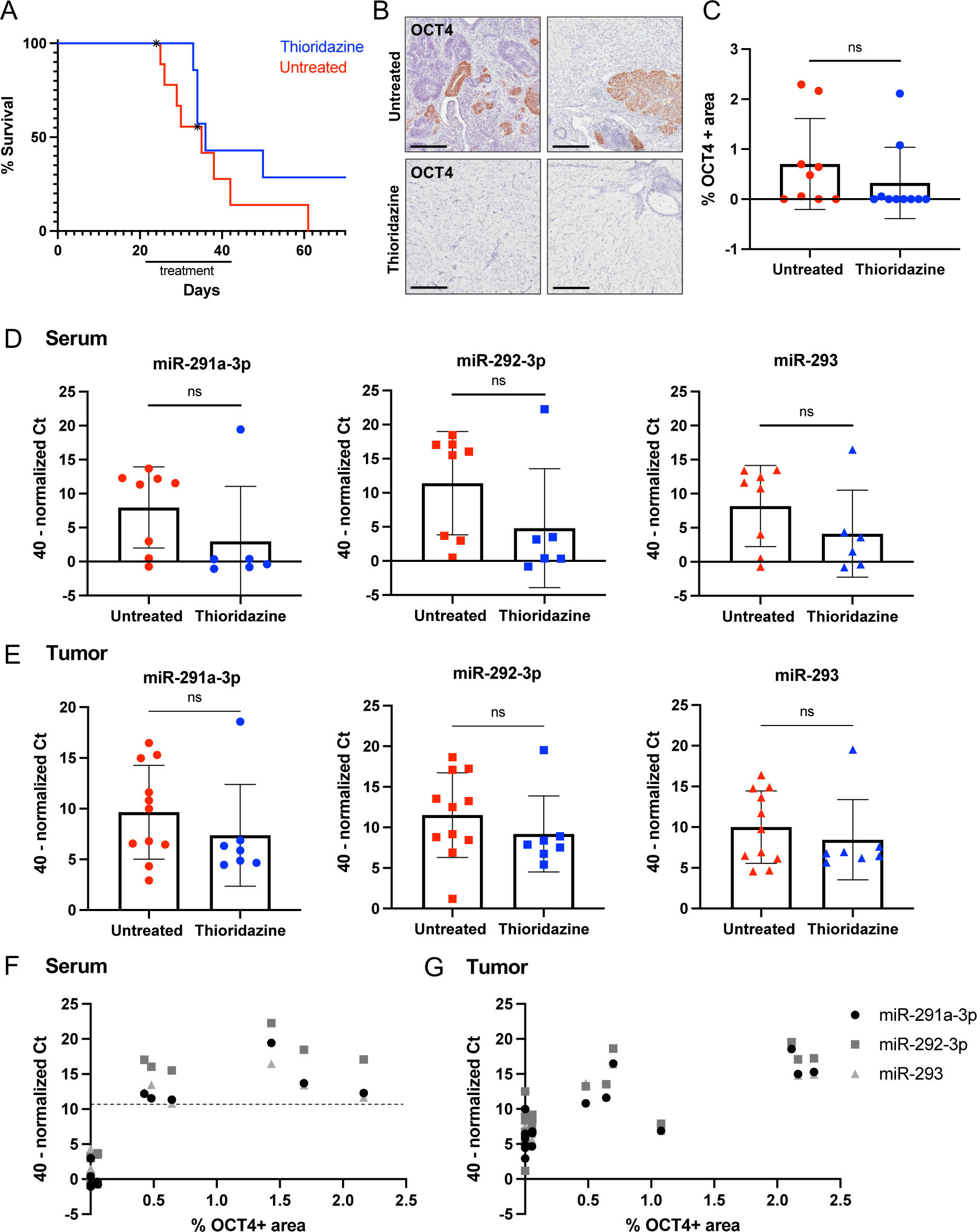
TR treatment reduces OCT4-positive EC cells in gPAK tumors, resulting in decreased serum miR-291-293 levels. (**A**) gPAK mice were treated with TR or vehicle control once every 3 days for 3 weeks starting at 21 days of age. Kaplan-Meier survival curve depicts the percentage of surviving control (n=9) and TR-treated mice (n=8) mice over time. One control mouse and one TR-treated mouse died during the study for reasons unrelated to the endpoint criteria and were therefore censored (as indicated by an asterisk). p = 0.2088 (Log-rank test). (**B**) Representative images of OCT4 IHC-stained tumors from control and TR treated mice. Scale bar = 200 μm. (**C**) Percentage of tumor cross-section (area) that was OCT4-positive as determined by IHC. One section per tumor was stained and quantified. (**D**) miR-291-293 expression in serum of control or TR-treated gPAK mice. (**E**) miR-291-293 expression in tumors of control or TR-treated gPAK mice. (**F & G**) Percent OCT4-positive area (data in C) versus miR-291-293 expression in (**F**) serum (data in D) or (**G**) tumor (data in E). In F, the percent OCT4-positive area per mouse for mice with two tumors was determined by calculating a weighted average of the percent OCT4-positive area per tumor section proportional to the whole tumor section area. Data represented as mean ± SD. ns = not significant (unpaired two-tailed t-test).

Knowing that gPAK tumors from TR-treated mice were in most cases depleted of EC, we next assessed miR-291-293 levels in the serum of TR-treated and control mice. Serum miR-291-293 levels were higher in control mice than in TR-treated mice, although the effect was not significant due to the one outlier mouse in the treatment group (Figure 4D). In fact, only the five untreated mice with tumors containing OCT4-positive cells had high levels of miR-291-293 in their serum. There was no significant difference in the levels of miR-291-293 in the tumor tissue from control and TR-treated mice (Figure 4E). These data suggest that elevated serum miR-291-293 levels, but not tumor miR-291-293 levels, indicate the presence of EC within a tumor. Furthermore, plotting the percent OCT4-positive area of a mouse’s tumor(s) against the serum miR-291-293 levels from that mouse revealed that miR-291-293 levels were above or below a theoretical threshold based on the presence or absence of EC within a tumor, although they do not perfectly correlate with the abundance of EC within a tumor (Figure 4F). Because there is more variation in tumor miR-291-293 levels, tumors with and without OCT4-positive cells cannot easily be distinguished based on tumor miR-291-293 levels alone (Figure 4G). These data demonstrate that serum miR-291-293 levels can be used to determine whether a mouse carries a tumor that contains pluripotent, malignant cells.

### Serum miR-291-293 levels can predict the presence of subclinical gPAK TGCTs

One of the most useful but unsubstantiated applications of the miR-371-373 biomarker is in screening seemingly healthy at-risk young men for the early detection of subclinical TGCTs. We therefore tested if serum miR-291-293 levels could be used to predict the presence of TGCTs in mice prior to clinical signs of the disease. Because TGCTs initiate around embryonic day 12.5 (E12.5) in the gPAK mouse model, we hypothesized that miR-291-293 could be detected in the serum of pregnant dams carrying a fetus with a TGCT. Timed matings were set up between breeders that would generate litters with both gPAK and control pups. Noon of the day on which a copulatory plug was found was defined as E0.5. Blood was collected from the dam at E18.5 and from all pups at 14 to 16-days of age (P14-16), before tumors were detectable by palpation. Mice were monitored for the development of testicular tumors and a terminal blood collection was performed at a humane endpoint or after six weeks of age, which is well beyond initial TGCT detection in gPAK mice (Figure 5A).^4^ Based on the established timing of TGCT development in gPAK mice, mice with testicular tumors at collection were assumed to have had tumors at P14-16 and E18.5.

**Figure 5.**
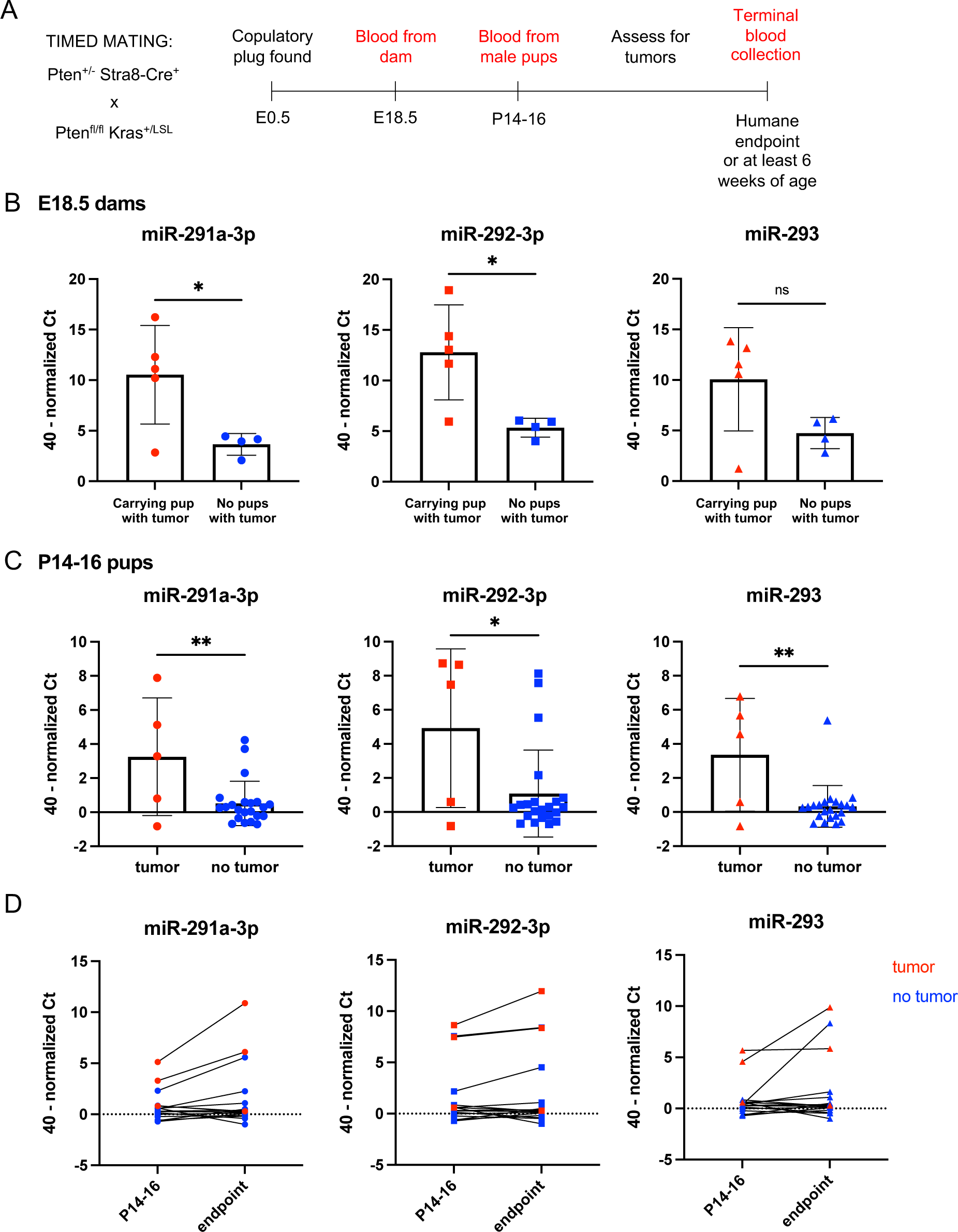
miR-291-293 expression predicts the presence of subclinical teratocarcinomas. (**A**) Schematic of experimental timeline for samples collected. (**B**) miR-291-293 expression in serum of pregnant dams (E18.5) carrying or not carrying a pup with a testicular tumor. (**C**) miR-291-293 expression in serum of 14-to 16-day old male pups that did or did not have a tumor. (**D**) miR-291-293 expression in serum of 14- to 16-day old mice and at endpoint (samples from the same mouse are connected by a line). ns = not significant, * p < 0.05, ** p< 0.01 (unpaired two-tailed t-test).

Analysis of serum from pregnant dams carrying a fetus that developed a TGCT showed significantly higher miR-291a-3p and miR-292-3p levels than that from pregnant dams that did not carry a fetus with a tumor (Figure 5B). Subsequent analysis of serum from individual pups at P14-16 revealed that serum miR-291-293 levels were significantly greater in pups that had a tumor (although at this time the tumor was not detectable by palpation) compared to control male pups without tumors (Figure 5C). Serum miRNA levels in tumor-bearing mice were generally lower at P14-16 than at humane endpoint when the tumor was clinically apparent; nevertheless, the trend shows that serum miR-291-293 levels at P14-16 correlated with serum miR-291-293 levels at endpoint (Figure 5D). Lower miR-291-293 expression at P14-16 may be due in part to much lower serum input volume, since only a small volume of blood could be collected from mice at this age. It is also worth noting that small sample volumes at this timepoint also were associated with lower assay specificity, as a few neonates without tumors showed similar miR-291-293 levels as the tumor-bearing littermates (Figure 5C). Together, these data demonstrate that serum miR-291-293 levels can be used to detect subclinical TGCTs, as early as during fetal development.

## DISCUSSION

Just as miR-371-373 are accurate biomarkers for malignant TGCTs in humans, we have found that miR-291-293 can serve as biomarkers of malignant TGCTs in mice, providing further evidence that the gPAK TGCT model closely resembles the human disease. Here, we have shown that both *in vitro* and *in vivo*, EC cells express and secrete miRNAs from the miR-290-295 cluster. Cultured EC cells derived from murine TGCTs express and secrete miR-291-293, and this expression is lost following differentiation by TR or SAL. This suggests that miR-290-295 expression is specific to the pluripotent state. It has previously been shown that the miR-290-295 cluster is a direct transcriptional target of OCT4, SOX2, and NANOG--all key regulators of pluripotency.^48^ Additionally, miR-290-295 expression enhances the efficiency of somatic cell reprogramming to iPS cells by OCT4, SOX2, and KLF4, but does not improve reprogramming efficiency in the presence of C-MYC because these miRNAs are downstream effectors of C-MYC.^49^ Therefore, pluripotency and miR-290-295 expression are intimately linked.

Not only are miR-290-295 expressed under the control of pluripotency-associated transcription factors, but they promote pluripotency themselves by regulating the expression of genes involved in multiple biological processes. Our results implicate miR-290-295 in regulating gene expression in cultured EC cells, as predicted targets of this miRNA cluster were upregulated in differentiated cells in which miR-290-295 expression is lost. In ES cells, members of the miR-290-295 cluster directly target and suppress multiple negative regulators of the G1/S transition, including *Cdkn1a* (encoding p21), *Rbl2*, and *Lats2*, thereby promoting proliferation.^23^ *Cdkn1a* and *Lats2* were significantly upregulated in TR3 and TR11 cells as compared to their parental EC cell lines, and *Rbl2* was significantly upregulated in TR11 cells. This suggests that these genes are being suppressed in EC cells to support progression through the G1/S transition and cell proliferation. After TR-mediated differentiation and loss of miR-290-295, expression of these cell cycle regulators increased. Consistent with this, TR-differentiated cells show moderately reduced proliferation compared to their respective parental EC cell line.^28^ The EC14 line is partially resistant to TR-mediated differentiation, with TR having less pronounced effects on proliferation, and accordingly the impacts of TR on miR-290-295 target gene expression were limited in this cell line.

miR-291-293 also were detected in the serum of mice bearing tumors containing EC cells. Allografts derived from cultured EC cells expressed miR-291-293 and secreted these miRNAs into the serum. Mice that received injections of differentiated cells, as well as wild-type 129S6 adult male mice, did not have elevated serum levels of miR-291-293. Interestingly, although EC-derived tumor tissue did have significantly higher expression of some of the miRNAs compared to normal testis tissue, the difference in expression between these tissue types was much smaller than the difference in miR-291-293 levels in the serum of the same mice. It has been previously shown that spermatogonia express miR-290-295, accounting for the expression in adult testis.^27^ It is possible that the spermatogonia that express miR-291-293 do not secrete it into circulation. Alternatively, the miRNAs could be secreted by spermatogonia but rapidly degraded while miRNAs secreted by EC cells into the serum are protected by extracellular vesicles (EVs). It is known that circulating miRNAs are often protected from endogenous RNases by association with protein complexes or encapsulation in EVs.^50^ A previous study showed that the human seminoma-derived cell line, TCam-2, secretes exosomes, which are small EVs, that contain miR-371a-3p.^51^ The functional roles for secreted miRNAs, including miR-371-373, in intercellular communication among TGCT cells or between TGCT cells and their microenvironment are currently unknown.

Consistent with previous findings that pure teratomas in humans do not express miR-371a-3p,^15^ we found that miR-291-293 expression was minimal in testicular teratoma tissue from *Dazl^-/-^*mice and undetectable in serum from teratoma-bearing mice. This suggests that miR-371-373 and miR-291-293 expression in humans and mice with TGCTs is specific to the malignant cells within the neoplasms. We also found that mammary adenocarcinomas from *MMTV-PyMT* mice did not express miR-291-293, confirming that expression of these miRNAs is not a common feature of malignant somatic cells.

Serum-based detection of EC cell markers in gPAK TGCTs could be an important tool in testing experimental therapeutics that aim to target EC cells, as these are the tumor-propagating cancer stem cells within these tumors. We previously showed that TR, an inducer of cell differentiation, significantly extended mouse survival in a human EC xenograft model.^28^ Here, we show that TR modestly increased the survival of tumor-bearing gPAK mice, and except for one non-responder, eliminated the OCT4-positive EC cells within tumors, suggesting that the increase in survival is due to the loss of the highly malignant EC compartment. In the gPAK model, mice develop spontaneous testicular tumors that are palpable around 21 days of age on average; however there is much variation in the age at which tumors are detectable.^4^ Because the size of tumors cannot be readily measured without advanced imaging tools when they are within the abdominal cavity, we started TR administration at 21 days of age in all treated mice, a time point at which the malignancy can be quite advanced. The variation in tumor size and disease progression at the time of treatment may explain why we did not detect a significant increase in survival with TR treatment in this study.

Paralleling EC loss following TR treatment, serum miR-291-293 levels were significantly reduced in TR-treated mice except for one non-responsive outlier. miR-291-293 expression was not significantly reduced in tumor tissue from TR-treated mice compared to that from control mice. This suggests that only tumors with an EC component secrete miR-291-293 into the serum at high levels. It is possible that low levels of miR-291-293 expression in tumor tissue from both control and TR-treated mice is from normal germ cells; however, the gPAK TGCTs usually lack all normal testicular architecture. Correlating the OCT4-positive area of a tumor with the miR-291-293 levels in the serum of the mouse bearing that tumor or in the tumor tissue itself, revealed that there is a clear diagnostic threshold of serum miR-291-293 abundance, above which it is predictive of the presence of an EC-containing tumor.

One of the most exciting potential applications of the miR-371a-3p biomarker in humans is as a screening tool to identify individuals with GCNIS lesions or localized TGCTs. If TGCTs are detected at an early enough stage, individuals can undergo surgery and active surveillance without the need for adjuvant chemotherapy or radiation^2^, highlighting the value of screening adolescent and young adult men in high-risk groups for GCNIS or otherwise undetectable TGCT lesions using the serum miR-371a-3p biomarker. However, in one study, miR-371a-3p levels were detectably increased in only 52% of patients with GCNIS lesions.^52^ Here, we found that miR-291-293 expression significantly increased in P14-16 mice with TGCTs that were not yet clinically detectable (Figure 5C). However, a larger sample size is needed to determine the sensitivity of this assay in detecting subclinical disease. Additionally, possibly due to the very small sample volume that can be collected from mice at this young age, miR-291-293 levels appeared to be increased in a few mice that did not have a tumor, reflecting potential specificity limitations of the assay in this specific context. It is unknown whether gPAK TGCTs have a GCNIS-like state of dormancy, so the subclinical detection of gPAK TGCTs may not accurately reflect detection of human GCNIS lesions, but rather early malignant TGCTs. It is possible that by using this mouse model, we could also discover ways to increase sensitivity in detecting GCNIS lesions. For example, if miR-291-293 are shed by EC cells in EVs, EVs could be isolated from human serum to enrich for miR-371a-3p. Improving the sensitivity of this assay may also allow for earlier detection of relapsing disease after treatment.

Since TGCTs initiate during embryogenesis in the gPAK mouse model, as they do in humans, we sought to determine if miR-291-293 could be detected in the serum of pregnant dams carrying a fetus that has initiated TGCT development. Indeed, maternal serum had significantly higher levels of miR-291a-3p and miR-292-3p when carrying a fetus with a TGCT compared to pregnant dams with no tumor-bearing pups. In only one out of five serum samples from dams carrying tumor-bearing pups did the assay fail to detect increased levels of these miRNAs. Additionally, none of the pregnant dams that were not carrying a tumor-bearing mouse had elevated levels of miR-291-293 (no false positives), which suggests that screening pregnant dams has even better specificity than screening P14-16 mice. It is possible that these very early TGCT lesions at E18.5 are more undifferentiated than they are at P14-16, and therefore express miR-291-293 at greater levels. Therefore, screening pregnant women whose sons would be at high risk for developing a TGCT (for example, due to a family history of TGCTs)^53^, may be more successful than the attempts made to screen men for GCNIS. This approach also might be informative for non-testicular malignant GCT diagnosed at early age as well.^54^ One caveat is that miR-371-373 are thought to be expressed by the placenta, as they are expressed by some trophoblast cell lines.^55^ This may obscure the miR-371-373 expression from early TGCTs developing in utero, but it is probable that TGCT expression of these miRNAs would far exceed the basal expression by the placenta, still making the assay diagnostically useful.

Together, these results further validate the gPAK mouse model as being representative of human TGCTs, highlighting its translational potential. Like malignant TGCTs in humans, gPAK TGCTs arise during embryonic development, express pluripotency markers, are sensitive to chemotherapy, and express mouse homologs of human miR-371-373. The finding that miR-291-293 can be used as serum biomarkers of malignant TGCTs in mice may facilitate *in vivo* drug screening to discover new therapies for TGCT treatment, particularly those aimed at targeting EC. Additionally, this mouse model can be utilized to better understand how miR-290-295 and miR-371-373 may contribute to TGCT pathogenesis. Finally, this model demonstrates the promise for future studies aimed at refining the miR-371-373 assay into a screening tool for malignant TGCT early detection.

## METHODS

### Cell culture

Murine EC cell lines were cultured and differentiated as previously described.^28^ For miRNA detection, cells at similar passage were plated, media was changed the following day, and 24 hours later the conditioned media was collected and centrifuged at 1500 rpm for 5 minutes to remove any cellular debris. The cells were pelleted and both the cells and conditioned media were stored at -80 °C prior to miRNA expression analysis.

### RNA sequencing and data analysis

RNA sequencing of EC3 and TR3 cells (*Pten^−/−^ Kras^+/+^ Stra8-Cre^Tg^ OCT4-gfp^Tg^*) was performed as previously described.^28^ RNA sequencing of EC11 and TR11 (*Pten^−/−^ Kras^+/+^ Stra8-Cre^Tg^*) and EC14 and TR14 cells (*Pten^−/−^ Kras^+/G12D^ Stra8-Cre^Tg^ OCT4-gfp^Tg^*) was performed in a previous study.^28^ Raw, un-normalized counts were filtered for expressed genes, defined by a minimum count value of 100 in at least one group. Standard differential analysis was done on the filtered genes using DESeq2 (v1.30.1)^56^ in R (v4.0.2) with R studio. Results for each cell line were extracted by comparing the EC cell line to its matched TR-differentiated derivative, using a false discovery rate cutoff of 0.05. Data were visualized using the *ggplot2* (v3.3.6), *dplyr* (v1.0.8), and *ggrepel* (v0.9.1) packages in R studio. *Gene Set Enrichment Analysis.* DESeq2 normalized count values per gene were averaged across three independent replicates per group, and log2 fold change values were calculated per gene to compare each TR-differentiated cell line to its parental EC cell line. Data were filtered to include only expressed genes, as defined by a minimum normalized count value of 100 in at least one group. Gene lists pre-ranked by log2 fold change were then created for each of the three comparisons. Pre-ranked gene lists and miRDB gene sets were used for Gene Set Enrichment Analysis, which was performed with 1000 permutations and a classic enrichment statistic.^29–31^

### Mouse strains and husbandry

All animals included in this study were handled in accordance with federal and institutional guidelines, under Institutional Animal Care and Use Committee-approved protocols. Mice were housed in facilities accredited by the Association for the Assessment and Accreditation of Laboratory Animal Care International and were cared for in compliance with the Guide for the Care and Use of Laboratory Animals.^57^ gPAK mice were generated as previously described.^4,58^ 129S6 and *MMTV-Pymt* mice were bred and maintained at Cornell University. *Dazl^-/-^* (*Dazl-1L*) mice were generated as previously described^47^ and backcrossed to the 129S4 genetic background.

### Genotyping

DNA was extracted from mouse tail snips and genotyped by PCR using primers specific for *Pten* and *Kras* alleles and the *Stra8-Cre* transgene as previously described.^58^ *Dazl^-/-^*mice were genotyped as previously described.^47^

### Subcutaneous cell transplantations

Tumor tissue derived from subcutaneous transplantation of cultured EC and differentiated cells into immunocompromised mice as well as serum from those mice were obtained from an experiment reported in a previous study.^28^

### Thioridazine treatment of gPAK mice

At 21 days of age, gPAK mice were randomly assigned to control or TR treatment groups. TR-treated mice were injected intraperitoneally with 25 mg/kg thioridazine hydrochloride (Sigma) dissolved in sterile 0.9% NaCl and sterile filtered, every 3 days for 3 weeks for 7 doses total or until humane endpoint criteria were met. Control mice received an equal volume of sterile 0.9% NaCl in the same manner. All mice were monitored regularly and euthanized by CO_2_ asphyxiation upon meeting humane endpoint criteria (loss of over 20% body weight, tumor greater than 2 cm in diameter, severe abdominal distension, displaying signs of pain such as hunching and piloerection) or after reaching 70 days of age. TR-treated or control mice that did not develop testicular tumors were excluded from the study (about 25% of gPAK mice do not develop tumors).^4^

### Serum and tissue collection

Blood was collected from pregnant females and neonates via the facial vein. For blood collection at endpoint, blood was obtained via intracardiac puncture immediately following euthanasia. Blood was allowed to clot at room temperature, then centrifuged at 5500 rpm for 10 minutes to separate serum from cellular components. Serum was transferred to a new tube and stored at -80 prior to miRNA expression analysis. Immediately after blood collection, tissue was dissected and flash frozen in liquid nitrogen, then stored at -80 prior to miRNA expression analysis.

### Immunohistochemistry and quantification

OCT4 immunohistochemistry was performed on one tissue section per tumor as previously described.^28^ OCT4-positive area per section was quantified using QuPath by manually selecting and annotating the areas of OCT4-postive staining and dividing the total OCT4-positive area by the total area of the entire tumor section.

### RNA isolation from serum and conditioned media

RNA from 100 µL of serum (35 µL for samples from pregnant dams and 15 µL for samples from P14-16 pups) or 200 µl conditioned media was isolated using the miRNeasy Serum/Plasma Advanced Kit (Qiagen) according to the manufacturer’s instructions. To increase RNA yield, MS2 carrier RNA (Roche) was added to a final concentration of 1.25 µg/mL. During lysis of the samples, a non-mammalian spike-in external control, cel-miR39 (5.6×10^8^ copies), was added to each sample in order to monitor RNA recovery. RNA was eluted from columns with 50 µL of nuclease-free water.

### RNA isolation from cells and tissues

Total RNA was extracted from tumor, testis, or spleen tissue or cell pellets using TRIzol Reagent (ThermoFisher) according to the manufacturer’s instruction. RNA quantity and quality were assessed using the Nanodrop One (Isogen Lifescience) and Qubit 4 fluorometer (ThermoFisher).

### miRNA expression analysis via TaqMan MicroRNA Assay

5 µL of purified RNA from serum or conditioned media samples or 10 ng of total RNA from tissue or cells was reversed transcribed using the TaqMan MicroRNA Reverse Transcription Kit (ThermoFisher) and the target-specific RT primer from the cel-miR39-3p (ID 000200), hsa-miR30b-5p (ID 000602), mmu-miR291a-3p (ID 002592), mmu-miR292-3p (ID 002593), and mmu-miR293-3p (ID 001794) TaqMan MicroRNA Assays (ThermoFisher). The final volume of 15 µL for each reaction underwent reverse transcription using a BioRad T100 Thermal Cycler at 16 °C for 30 min, 42 °C for 30 min, followed by a final step of 85 °C for 5 min. For the final TaqMan PCR, for each of the aforementioned TaqMan MicroRNA Assays, 1.5 µL of the cDNA product was added to 10 µL of 2X TaqMan Fast Advanced Master Mix (ThermoFisher) and 1 µL of 20X TaqMan Assay (including the TaqMan probe and PCR primer set) and brought up to a final reaction volume of 20 µL. All reactions were performed in duplicate. Ct values were measured on a QuantStudio 12K Flex device. “Undetected” calls were assigned a Ct value of 40 (maximum number of cycles) so that quantification was possible. The cel-miR39-3p assay was performed for serum and conditioned media samples as a quality control check to monitor RNA recovery. For normalization, endogenous reference miR-30b-5p was used. Targets were corrected for average miR-30b-5p levels across biological replicates to correct for deviations in the endogenous levels of miR-30b-5p. 40 minus delta Ct transformation was used to represent miRNA expression levels.

### Statistical analysis and visualization

Except for analysis of RNA sequencing data, all other statistical analyses were performed using GraphPad Prism. Data were visualized using GraphPad Prism.

## ACKNOWLEDGEMENTS

A.R.L. was funded by NCI F30 CA247458. D.M.T., A.J.M.G. and L.H.J.L. were financially supported by the Princess Máxima Center for Pediatric Oncology through KiKa funding. The authors would like to thank Jen Grenier from the Transcriptional Regulation and Expression facility at Cornell University for RNA sequencing and assistance with data analysis, Thao. T. Pham for genotyping of *Dazl* mice, and Praveen Sethupathy and Andrew Grimson for helpful discussions.

