## Supplemental Materials for "Analysis of a mouse germ cell tumor model establishes pluripotency-associated miRNAs as conserved serum biomarkers for germ cell cancer detection"

**Supplemental Table 1.** Genes that were experimentally proven to be direct targets of the miR-290-295 cluster and their associated publication.

| Mouse gene name | Reference |
|---|---|
| ***Tgfbr2*** | Guo et al. 2015[32] |
| ***Gsk3b*** | Guo et al. 2015[32] |
| ***Arhgef3*** | Guo et al. 2015[32] |
| ***Fndc3a*** | Guo et al. 2015[32] |
| ***Vim*** | Guo et al. 2015[32] |
| ***Cdkn1a*** | Wang et al. 2008[23] |
| ***Rbl2*** | Wang et al. 2008[23] |
| ***Lats2*** | Wang et al. 2008[23] |
| ***Casp2*** | Zheng et al. 2011[24] |
| ***Ei24*** | Zheng et al. 2011[24] |
| ***Wee1*** | Lichner et al. 2011[22] |
| ***Fbxl5*** | Lichner et al. 2011[22] |
| ***Dkk1*** | Zovoilis et al. 2009[33] |
| ***Rela*** | Lüningschrör et al. 2012[34] |
| ***Pax6*** | Kaspi et al. 2013[35] |
| ***Ccnd1*** | Gong et al. 2017[36] |
| ***Arid4b*** | Goldberger et al. 2013[37] |
| ***Mbd2*** | Cao et al. 2015[38] |
| ***Ash1l*** | Kanellopoulou et al. 2015[39] |
| ***Tfap4*** | Schaefer et al. 2022[40] |
| ***Runx2*** | Akshaya et al. 2022[41] |
| ***Pfn2*** | Sangokoya and Blelloch 2020[42] |
| ***Atg5*** | Lu et al. 2019[43] |
| ***Becn1*** | Lu et al. 2019[43] |
| ***Mbnl1*** | Wu et al. 2018[44] |
| ***Mbnl2*** | Wu et al. 2018[44] |

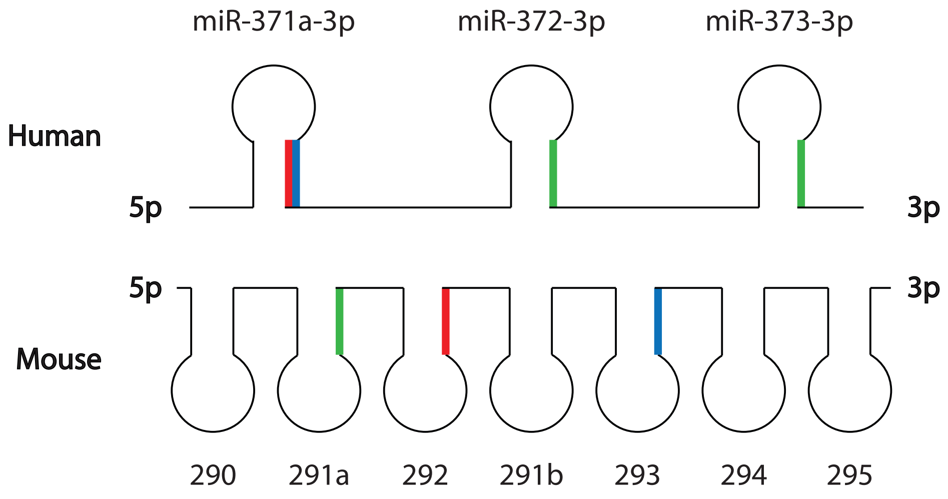


**Supplemental Figure 1.** Homology of the miR-371-373/miR-290-295 cluster family. Mouse miR-291a-3p is homologous to human miR-372-3p and miR-373-3p, whereas miR-292-3p and miR-293 are homologous to different isoforms of human miR-371a-3p.

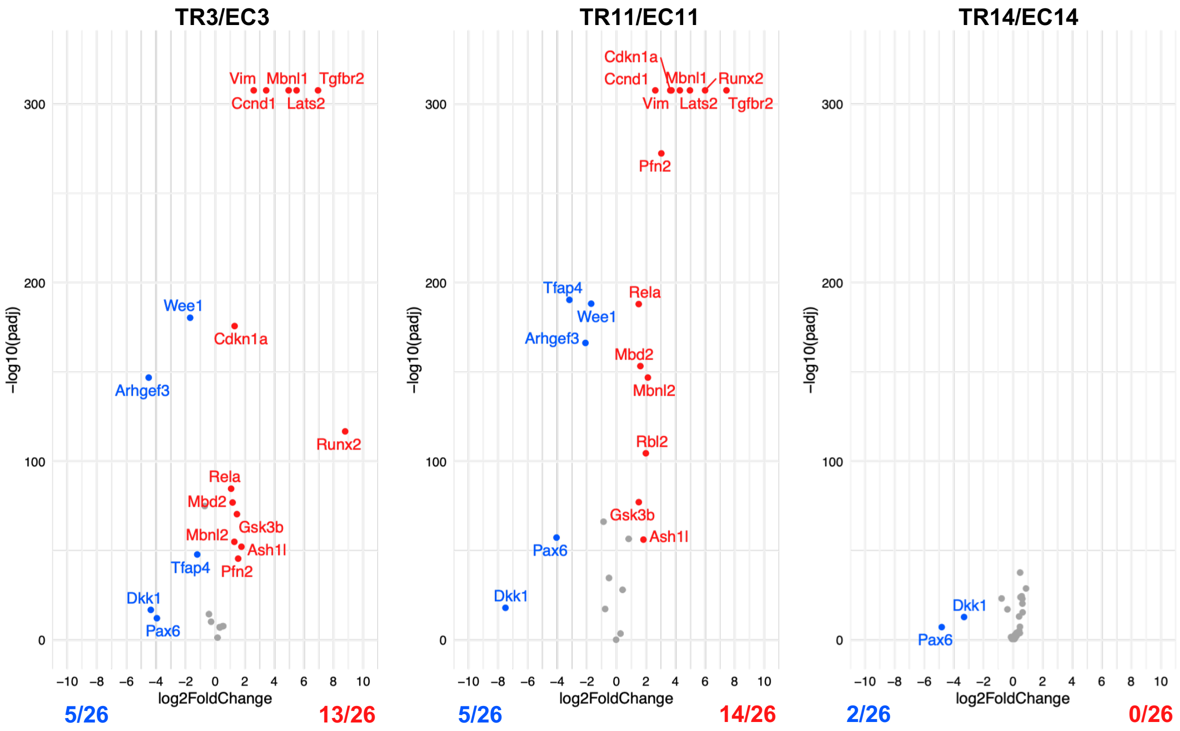


**Supplemental Figure 2.** Volcano plots of 26 experimentally proven direct target genes of the miR-290-295 cluster (see Supplemental Table 1). Genes that are significantly upregulated in TR-differentiated cells are in red, genes that are significantly downregulated in TR-differentiated cells are in blue, and genes that are not significantly differentially expressed between the cells are in grey. Significance is defined as an adjusted p value (padj) $< 0.05$.
